# Personalization of mathematical models of human atrial action potential

**DOI:** 10.1101/2020.06.29.174870

**Authors:** Andrey V. Pikunov, Roman A. Syunyaev, Vanessa Steckmeister, Ingo Kutschka, Niels Voigt, Igor R. Efimov

## Abstract

Atrial cardiomyocytes demonstrate a wide spectrum of patient-specific, tissue-specific, and pathology-specific action potential (AP) phenotypes due to differences in protein expression and posttranslational modifications. Accurate simulation of the AP excitation and propagation in healthy or diseased atria requires a mathematical model capable of reproducing all the differences by parameter rescaling. In the present study, we have benchmarked two widely used electrophysiological models of the human atrium: the Maleckar and the Grandi models. In particular, patch-clamp AP recordings from human atrial myocytes were fitted by the genetic algorithm (GA) to test the models' versatility. We have shown that the Maleckar model results in a more accurate fitting of heart rate dependence of action potential duration (APD) and resting potential (RP). On the other hand, both models demonstrate the poor fitting of the plateau phase and spike-and-dome morphologies. We propose that modifications to L-type calcium current-voltage relationships are required to improve atrial models' fidelity.

## 1 Introduction

Atrial fibrillation (AF) is the most common chronic arrhythmia encountered in clinical practice [1]. Today, radio-frequency catheter ablation is considered to be one of the most effective treatment options, however, the successful outcome is reported to be from 56% to 89% by different clinical trials, moreover, post-operational complications are quite common [2]. On the other hand, several groups reported that personalized mathematical simulation of the action potential (AP) propagation in atria is capable to reproduce clinical measurements and consequently might be used to improve the outcome of the operation by choosing optimal ablation targets [3, 4]. One important challenge that remains in the field is inter-subject AP variability present in cardiac electrophysiological recordings [5]. Indeed, despite the variety of AP waveform morphologies recorded from atrial myocytes [6–12] both experimental and theoretical studies are mostly focusing on reporting the averaged characteristics of AP. To accurately describe the phenotypic variability, the baseline mathematical model should be able to reproduce these differences by the means of model parameter rescaling. While several groups benchmarked the existing human atrial models [5, 13] the question of whether these models are flexible enough to be personalized to a particular AP waveform remains to be answered.

Previously we have developed a genetic algorithm-based (GA-based) solution that was capable of precise personalization of the O’Hara-Rudy model [14] using optical AP recordings of the human left ventricular wedge preparations [15]. In the present preliminary research, we have utilized similar techniques to investigate the capability of the Maleckar [16] and the Grandi [17] atrial AP models to describe the variability of the AP morphology in healthy and diseased human atria.

## 2 Methods

### 2.1 Experimental data acquisition

#### Human tissue samples and myocyte isolation

Right atrial appendages were obtained from patients undergoing cardiac surgery. Experimental protocols were approved by the ethics committee of the University Medical Center Göttingen (No. 4/11/18). Each patient gave a written informed consent. The patient samples were grouped in four blinded sets according to the diagnosis: sinus rhythm (SR), postoperative AF, paroxysmal AF, and chronic AF (referred to as Groups 1 - 4 below). Excised right atrial appendages were subjected to a standard protocol for myocyte isolation [18]. Right atrial myocytes were suspended in EGTA-free storage solution for subsequent simultaneous measurement of membrane current/potential and intracellular Ca^2+^ concentration.

#### Measurements of cellular electrophysiology

Only rod-shaped myocytes with clear striations and defined margins were selected for measurements of [Ca^2+^]_i_ and cellular electrophysiology. Cells were loaded with 10 μmol/L Fluo-3-AM (Invitrogen; 10 min loading and 30 min de-esterification) for simultaneous measurements of action potentials (AP) and intracellular [Ca^2+^]_i_. APs were recorded using current-clamp configuration. To evoke APs, 1 ms current pulses of 1.5-2x threshold strength were applied. Sometimes RP was too low to generate a proper AP since Na-channels were inactivated. In such cases, the small artificial hyperpolarizing current was introduced leading lower RP. The pacing cycle length (PCL) was decreased in a stepwise manner, starting at 2064 ms and subsequently 1032, 516, 344, 258, 206, 172, 148, 129 ms. During experiments, myocytes were perfused with a bath solution at 37 °C containing (in mmol/L): CaCl_2_ 2, glucose 10, HEPES 10, KCl 4, MgCl_2_ 1, NaCl 140, probenecid 2; pH=7.35. The pipette solution contained (in mmol/L): EGTA 0.02, GTP-Tris 0.1, HEPES 10, K-aspartate 92, KCl 48, Mg-ATP 1, Na_2_-ATP 4, Fluo-3 pentapotassium salt 0.1; pH=7.2. About 750μl/50 ml KOH 1M was added to the pipette solution to adjust pH. For the bath solution pH adjustment, approximately 3 ml of NaOH 1M was added 1l Tyrode solution. Pipette resistances in the range of 5-7 MΩ were utilized. Intracellular Ca^2+^ measurements were not considered in the present study.

### 2.2 Computer simulations

The custom C++ code replicating the Maleckar [16] and the Grandi [17] models was used to simulate AP of human atrial myocytes. The parameter values were fitted via GA as described in [15]. Briefly, a population of models with random parameter values was generated. Each model was paced for 9 beats at each experimental PCL before the fitness function (RMSE) evaluation. Next, the model population was modified by mutation and crossover genetic operators, followed by tournament selection. The cycle was repeated for each new generation. AP waveforms at PCLs of 2064, 1032, 516, 344, 258, 206, and 172 ms were used as the input traces for GA runs. Although intracellular Na, K, and sarcoplasmic Ca concentrations were modified by genetic operators, Na and K concentrations were limited to narrow ranges corresponding to experimental pipette solutions ([7, 9] and [130, 150] mM, respectively). Two additional parameters were adjusted by GA: the amplitude of the stimulation current (I_st_) and the constant micropipette current distorting AP waveform (I_bl_). The latter accounts for possible experimental artifacts and represents a combination of the hyperpolarizing current introduced to control the RP level and the leak current. The ranges for I_st_ and I_bl_ were set to [−80, −20] pA/pF and [−1, 1] pA/pF, respectively. Multipliers for model ionic current magnitudes were limited to [0.001, 10] range.

The total number of organisms in a population was 320 with 24 elite organisms that were not modified by genetic operators. The number of generations in a GA run was set to 400. Simulations were performed on the cluster of The Moscow Institute of Physics and Technology (Intel Xeon Processor E5-2690, 2.90GHz) and the George Washington University’s Pegasus cluster (Intel Xeon Gold 6148 Processor, 3.70GHz). In the case of equal numbers of organisms and CPU cores, a single GA run took about 5 hours for the Maleckar model and 20 hours for the Grandi model.

## 3 Results

### 3.1 Experimental AP waveforms

Experimental AP waveforms and dependencies of APA, APD80, and RP on PCL for all 4 groups of cells are shown in Figure 1. We have observed a substantial difference in the AP morphologies not only across patients of the different groups but also among the patients of the same group. For example, Group 1 Cell 1 demonstrates a short ventricular-like plateau phase at +25 mV, while other cells from the same group have a more atrial-like triangulated AP waveform. Other groups of patients demonstrate both triangulated and spike-and-dome waveforms, that were previously characterized as typical to AF and SR, respectively [7, 9]. It should be noted that in some cases RP dependence on PCL is nonmonotonic (e.g. Cells 1 and 2 from Group 2) resulting in corresponding distortions of APA and APD curves. Such RP behavior could be explained by the effect of the I_bl_ described in the previous section‥

**Fig. 1.**
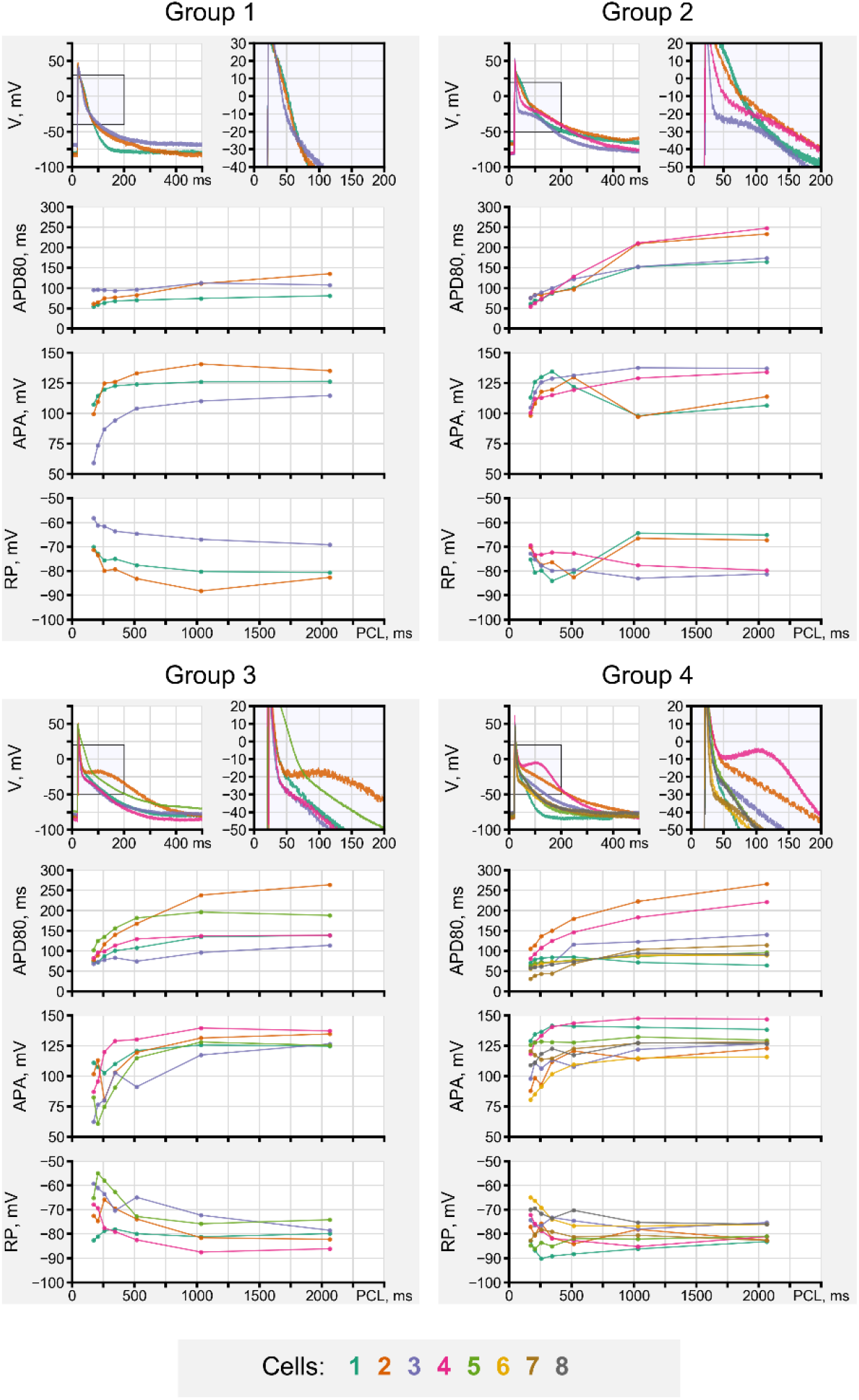
AP waveforms, APD80, APA, and RP for cells of 4 groups recorded in the patch-clamp experiment. AP waveforms are drawn at PCL 2064 ms. PCLs 2064, 1032, 516, 344, 258, 206, and 172 were used for APD80, APA, and RP curves.

### 3.2 Genetic algorithm runs

#### Model comparison

Output model RMSE scores for the Maleckar and the Grandi models were 3.5 (2.5 — 3.8) mV and 4.3 (3.3 — 4.8) mV, respectively (errors will be given hereinafter as a median followed by an interquartile range in brackets). Although the overall difference in RMSE scores was not significant, it should be noted that in every single case the Maleckar model reproduced the experimental AP with lesser RMSE than the Grandi model. On the other hand, GA output concentrations for the Grandi model were closer to the steady-state solution (data is not shown).

Representative fitting results for both models are shown in Figure 2: triangular waveform (Figure 2A) and spike-and-dome waveform (Figure 2B,C). For the cases shown in Figure 2A,B the fitting is relatively accurate for both models, however, the Maleckar model replicated overall AP morphology better, which is especially evident for larger PCLs. Figure 2C exemplifies the case when the Maleckar model failed to capture spike-and-dome morphology at large PCLs still being able to fit the APs at smaller PCLs, while the Grandi model did not reproduce the experiment within the whole range of the PCLs.

**Fig. 2.**
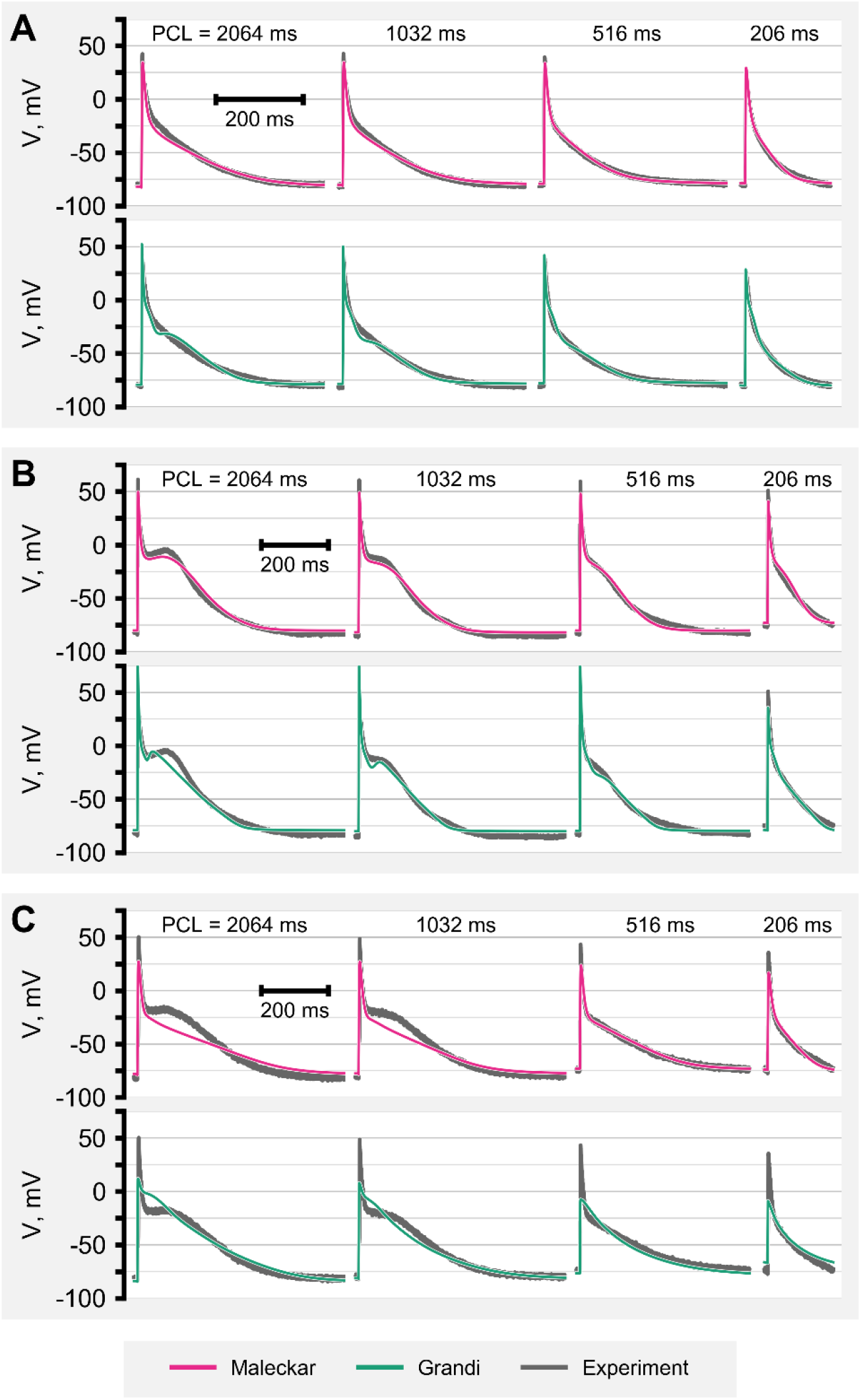
Comparison of the model and experimental AP waveforms. (A) Group 3 Cell 1. (B) Group 4 Cell 4. (C) Group 3 Cell 2.

Figure 3 compares the fitting errors for common electrophysiological markers (APD, APA, and RP). The errors of APD80 and APD60 of the Grandi model have a similar trend, varying from more positive values at PCLs less than 1032 ms to more negative at PCLs 1032 and 2064 ms. The difference between the two models is more prominent for APD60: the Grandi model interquartile range for the APD60 error was about twice bigger than the same value for the Maleckar model within all the PCLs (excluding PCL 1032 ms). The overall APA error for the Maleckar model equaled −8.6 (−15.1 — −2.3) mV and depended weakly on the PCL, while the Grandi model reproduced the APA with greater errors’ range at bigger PCLs (e.g. 0.9 (−17.4 — 3.8) mV for PCL 2064 ms). The RP error for the Grandi model monotonically depended on PCL and raised from −2.8 (−6.7 — −0.2) mV at PCL 172 ms to 1.7 (0.4 — 2.3) mV at PCL 2064 ms. Generally, the RP of the Maleckar model slightly outreached the experimental level (0.90 (−0.63 — 2.25) mV).

**Fig. 3.**
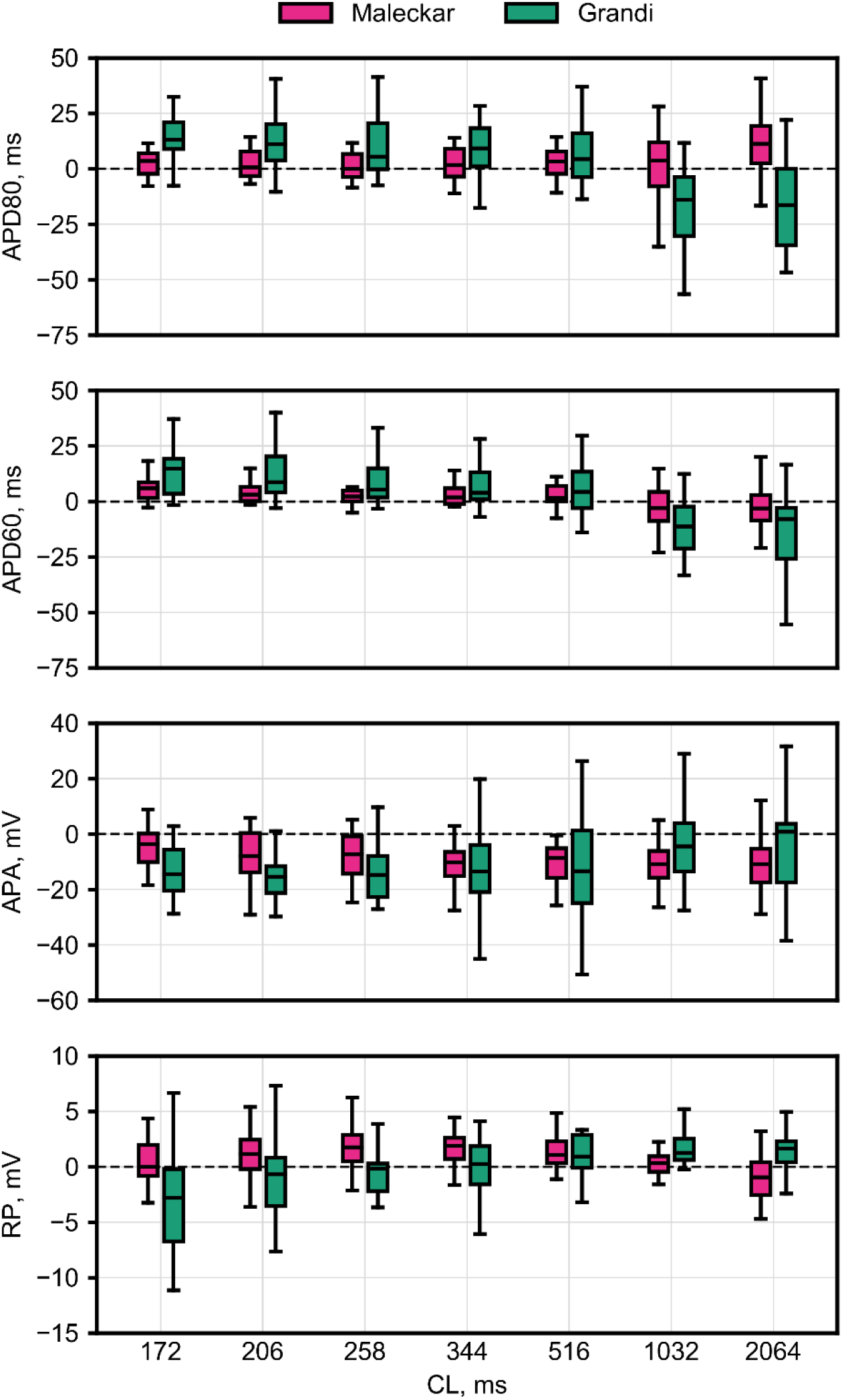
Errors in reproducing APD80, APD60, APA, and RP for the Maleckar and the Grandi models. Outliers are not shown.

Relying on the foregoing comparison, we considered utilizing the Maleckar model for further analysis, since this model was capable of reproducing APs more accurately as compared with the Grandi model.

#### Analysis of GA runs

The GA output parameters of the Maleckar model are demonstrated in Figure 4. In case of I_Na_, I_Kur_, and I_K1_, relatively narrow conductivity variation was required to explain the diversity of waveforms that we have observed in the experiment (median (interquartile range): 0.71 (0.63 — 0.83), 0.67 (0.54 — 0.89), and 0.84 (0.72 — 1.17) for I_Na_, I_Kur_, and I_K1_, respectively). Similar parameters’ ranges were reported previously by [5]. On the other hand, several ionic currents required extreme changes to model parameters to replicate the experimental AP waveform. In particular, I_Ca,L_ conductivity was less than 0.1 of the original Maleckar model conductivity, in 12 cases out of 20. On the contrary, I_Ks_ and I_Kr_ increase was above 500% of the original model value in 11 cases out of 20 for each of both currents. As discussed below, we hypothesize that these discrepancies in model parameters might arise because of inaccuracies in ionic currents formulations.

**Fig. 4.**
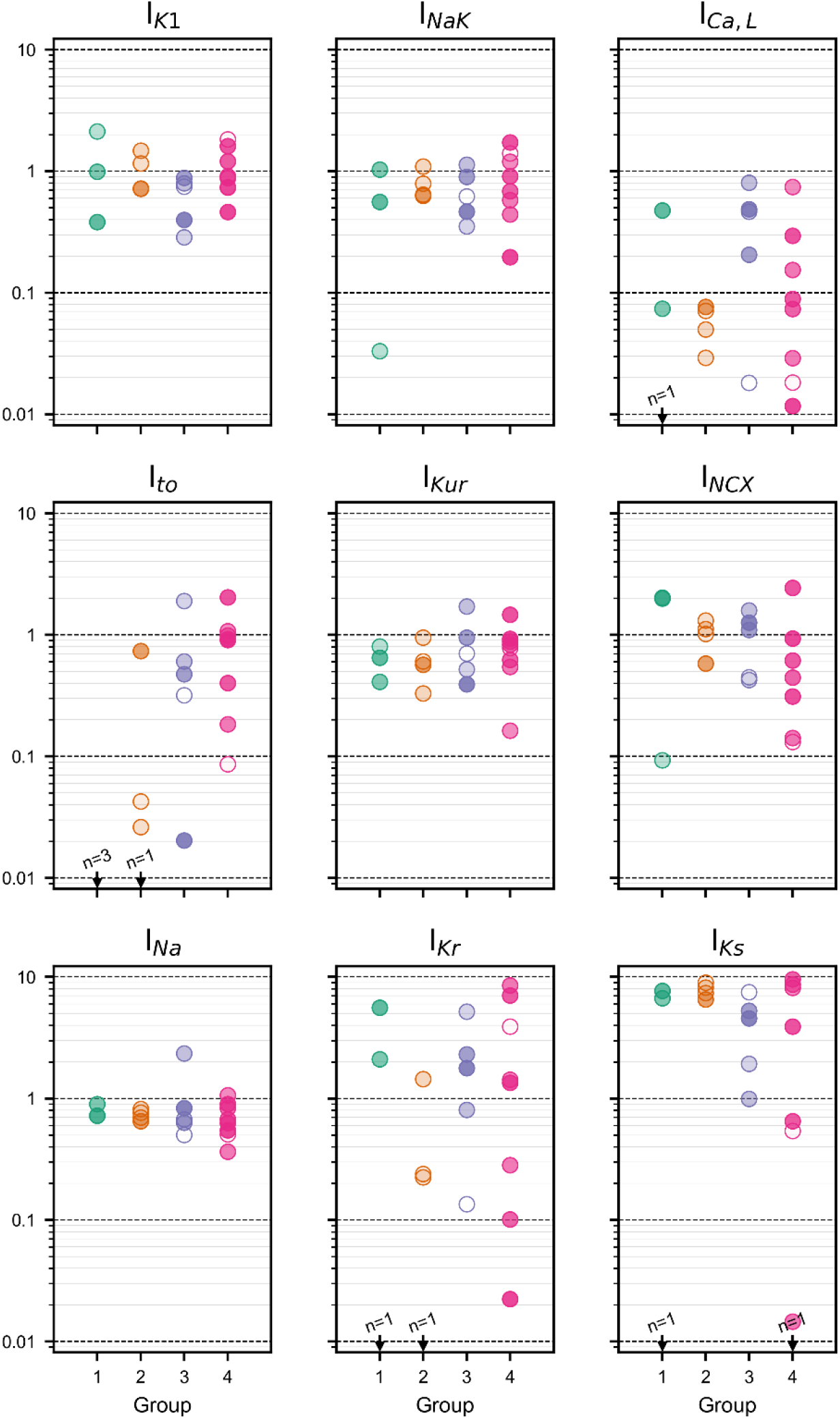
GA output parameters for the Maleckar model. Transparency of marker correlates with the fitting accuracy: bigger RMSE, more transparent the marker. Magnitudes lower 0.01 are indicated by arrows with a number of precedents.

## 4 Discussion

Long-term persistent AF is associated with extensive atrial remodeling and, in particular, with changes in ionic channels conductivities [19]. These changes shorten the APD [7–10, 20], causing a decrease in effective reentrant wavelength, which affects the sustainability of arrhythmia [21]. Therefore, quantification of the gradual remodeling associated with a transition from healthy to diseased phenotype is important.

Previously, using human ventricular AP recordings, we have demonstrated that GA accurately determines conductivities of high-amplitude ionic currents [15]. Applying GA to the atrial recordings from patients on different stages of AF development, we pursued twofold goals. First, to quantify the aforementioned remodeling of atrial ionic currents. Second, to personalize single-cell models for further simulations with real patient's heart geometry, focused on an investigation of the arrhythmogenesis during AF. The obvious limitation of personalization technique is that the current-voltage relationships and kinetics of ionic currents in the baseline model must correctly reproduce the experimental data. Otherwise, inaccurate formulations of an ionic current essentially reduce the identifiability of model parameters: the best waveform fitting results from an implausible combination of ionic currents conductivities which compensate for inaccuracies in current-voltage relationships.

We compared the two most widely used atrial models in terms of the ability to describe a broad spectrum of healthy and diseased atrial AP morphologies. While we observed that the Maleckar model, in general, resulted in a better fitting in comparison with the Grandi model (Figures 2 and 3), both of them failed to replicate all the variety of AP waveforms. This was especially evident in cases of a spike-and-dome morphology with a “deep notch” following an initial depolarization spike (Figure 2C). On the other hand, we have observed very low GA output I_Ca,L_ conductivity for the Maleckar model as shown in Figure 4. Taken together, denoted facts might indicate a substantial error in the formulation of I_Ca,L_.

Altered expression of the auxiliary subunits of the L-type calcium channel (namely α1C\α2δ1 and α1C\α2δ2, encoded by CACNA2D1 and CACNA2D2 genes, respectively) was previously reported for patients with AF [22]. According to studies of the differential expression, the CACNA2D1 expression is higher in ventricles, while the CACNA2D2 is typical for atria [23]. As shown in Figure 5C, current-voltage relationships of atrial [24] and ventricular [25] I_Ca,L_ differ. Primarily, this difference is caused by the corresponding activation curves (Figure 5D). We hypothesize that I_Ca,L_ in atria is presented as a composition of several subpopulations of ionic channels formed by different subunits and, consequently, characterized by different current-voltage relationships. The balance between these subpopulations may change during AF development [22]. Given the different current-voltage relationships of corresponding ionic currents, the alterations in the expression profile can not be replicated by a model with a single I_Ca,L_ subpopulation.

**Fig. 5.**
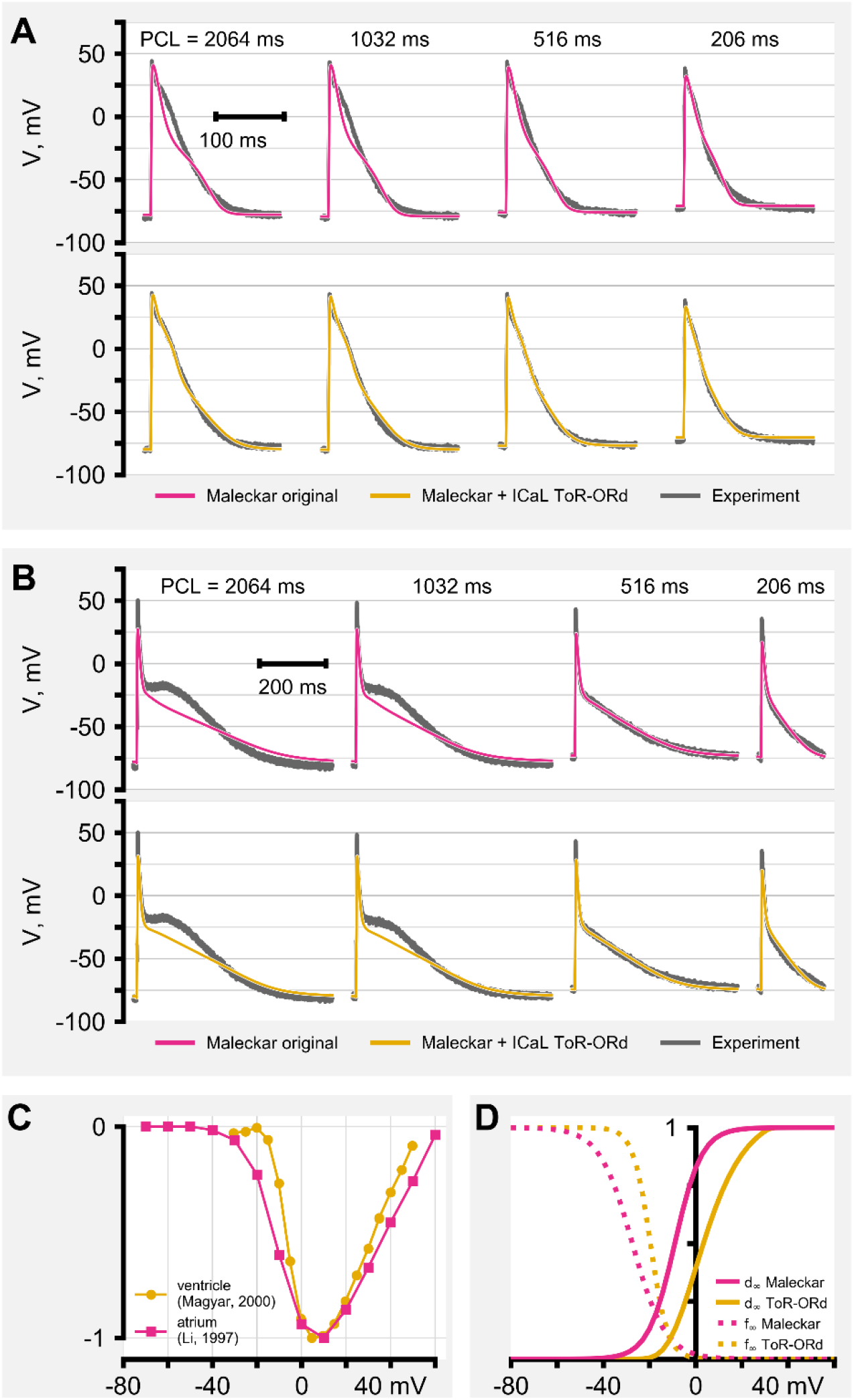
Model fitting for Group 1 Cell 1(A) and Group 3 Cell 2 (B). (C) Current-voltage relationships of atrial and ventricular I_Ca,L_. Adapted from [24] and [25]. (D) I_Ca,L_ activation curves for the Maleckar and ToR-ORd models‥

In line with the proposed hypothesis, we have introduced an additional ventricular subpopulation of I_Ca,L_ to the Maleckar model. The I_Ca,L_ formulation was adapted from [26]. The representative results of GA runs with the hybrid model are presented in Figure 5. In general, this modification indeed resulted in a more accurate fitting in comparison with the original model. For example, we have observed a 40% decrease in the fitting error (RMSE) for Group 1 Cell 1 (Figure 5A). Model AP approximated experimental waveforms much better in the region from −25 to 25 mV. However, the introduction of the ventricular I_Ca,L_ to the Maleckar model did not improve the fitting of the spike-and-dome morphology (Figure 5B), although RMSE was 10% lower. The reason for such inefficacy can be explained via I_Ca,L_ current-voltage relationships and activation curves (Figure 5C, D). The notch following the spike repolarizes the cell as low as −25 mV, while both subpopulations of I_Ca,L_ are essentially deactivated at this voltage level. We conclude that the absence of substantial depolarization current at −25 mV in the model prohibits the following “dome” observed in the experiment. It should be reiterated here that the presence of two subpopulations of I_Ca,L_ most probably affects the voltage-clamp experiments underlying the atrial models. Given that α2δ1 subunit expression is not exclusive to ventricles, actual atrial-specific α2δ2 activation might be shifted even further to negative voltages than the one measured in experiments. We also suspect a change of the balance between these two isoforms to result in a proarrhythmic substrate in the case of the long-term persistent AF. As presented in dynamic patch-clamp experiments [27], the alterations in the atrial I_Ca,L_ current-voltage relationships might result in early afterdepolarizations and trigger the arrhythmia. Further theoretical and experimental evidence is required to test these assumptions that we will address in the future studies.

## Fundings

This work was supported by grants from the Deutsche Forschungsgemeinschaft (DFG) to NV (VO 1568/3-1, IRTG1816, and SFB1002 project A13), from the Else-Kröner-Fresenius Foundation to NV (EKFS 2016_A20), from NIH/NHLBI (U01 HL141074) to IE, by Russian Foundation for Basic Research grants 18-07-01480, 19-29-04111, 18-00-01524 to RS.

## Acknowledgements

The authors thank Ines Müller and Stefanie Kestel for excellent technical assistance and the cardiac surgeons of the Department of Thoracic and Cardiovascular Surgery, University Medical Center Göttingen for kindly providing human atrial tissue samples.

## Notes

### Competing Interest Statement

The authors have declared no competing interest.

